# DNA-Dependent Protein Kinase Inhibitors PI-103 and Samotolisib Augment CRISPR/Cas9 Knockin Efficiency in Human T Cells

**DOI:** 10.1101/2024.10.18.618985

**Authors:** Emina Džafo, Morteza Hafezi, Greta Maria Paola Giordano Attianese, Patrick Reichenbach, Stephane Grillet, Kirsten Scholten, George Coukos, Melita Irving, Bernhard Gentner

## Abstract

The adoptive cell transfer of *ex vivo* expanded tumor infiltrating lymphocytes (i.e., TIL therapy) is a promising clinical strategy and recently FDA approved for melanoma but has major limitations including that not all tumors are inflamed. Moreover, tumor-specific clones can be rare and in an exhausted state due to the suppressive tumor microenvironment. These obstacles can be overcome by engineering autologous peripheral blood T cells with pre-selected T cell receptors (TCRs) by viral vector-mediated gene insertion. While viral transduction is highly efficient, the insertional site is not specific and persistence of the T cells is oftentimes limited. In contrast, site-specific integration of the TCR into the TCR α chain (*TRAC*) locus by CRISPR/Cas9 has been shown to enable more consistent and physiological levels of exogenous TCR expression coupled with superior persistence and tumor control in preclinical studies. Here, we sought to improve the efficiency of CRISPR/Cas9 mediated TCR knockin (KI) into the *TRAC* locus of primary human T cells. In addition to the previously reported DNA-dependent protein kinase inhibitor M3814, we demonstrate that PI-103 and samotolisib markedly increase KI efficiency in a process that is GMP-compatible, while CC-115 had a variable effect. Importantly, PI-103 and samotolisib do not negatively impact cell viability, fold-expansion nor T cell phenotype and we conclude that they are suitable for the generation of gene-modified T cells for clinical use.

## 2. Introduction

Adoptive cell therapy (ACT) is a promising strategy for the treatment of advanced cancer patients. In the setting of solid tumors, the administration of *ex vivo* expanded TILs enriched from patient biopsies has been shown to mediate durable regression of metastatic melanoma in some patients (1-5). Major limitations of this approach, however, are that not all tumors are inflamed, and the tumor-specific clones can be rare, in an exhausted state, and difficult to expand. Alternatively, polyclonal T cells derived from the peripheral blood mononuclear cells (PBMCs) and of a defined phenotype can be genetically engineered with a T cell receptor (TCR) or a chimeric antigen receptor (CAR) directed against tumor antigen enabling treatment with a defined cellular product (6).

Most engineered T cells used in the clinic have been generated with viral vectors (7). Although both lentivirus and gamma-retrovirus have a strong safety record, there are inherent risks due to semi-random genomic integration and the potential for oncogenesis (*4-7*). Moreover, reliance upon strong and constitutively active promoters may contribute to premature exhaustion of the T cells. Non-viral editing by CRISPR/Cas9 offers unprecedent precision with the possibility of expressing TCRs within the *TRAC* or TCR β chain (*TRBC*) locus for more natural and consistent expression levels. Early phase I clinical studies have demonstrated feasibility and promise of CRISPR/Cas9 based TCR-engineered T cells, including with neo-antigen specific TCRs (8-14).

Double-stranded breaks caused by Cas9 are repaired through two major repair mechanisms, non-homologous end joining (NHEJ) and to a lesser extent by homology-directed repair (HDR) which is the pathway needed for the KI of genes (15). The pharmacological inhibition or deletion of key NHEJ factors such as the DNA-dependent protein kinase (DNA-PK) can tip the scale towards HDR and allow the genomic integration of a donor template DNA sequence (16-22). Certain DNA-PK inhibitors such as M3814 favored HDR which led to an increased KI efficiency in engineered T cells (23,24).

Here, we compared the capacity of different DNA-PK inhibitors to increase CRISPR/Cas9 KI efficiency. In addition to the previously reported small molecule M3814, we present alternative compounds, PI-103 and samotolisib, which markedly increase the KI efficiency, while CC-115 showed variable results. The compounds were integrated into a good manufacturing practice (GMP)-compatible procedure and hold potential for more efficient generation of TCR KI T cells for clinical translation.

## 3. Results

With the goal of establishing an efficient CRISPR/Cas9 mediated TCR KI strategy that could ultimately be used for GMP-grade production of TCR-engineered T cells, we first gene-edited human T cells to express the murine marker Thy1.1 (CD90.1) as a reporter gene at the *TRAC* locus. Our preliminary strategy included the co-electroporation of Cas9 protein preincubated with guide (g)RNA described in Schober et. al. (9) targeting the *TRAC* locus to generate ribonucleoprotein (RNP), along with the Thy1.1 gene-insert in plasmid format. In brief, T cells were electroporated on day 2 post-activation and treated with four different DNA-PK inhibitors (M3814, CC-115, PI-103 and samotolisib) under standard culture conditions in serum-containing medium (Figure 1A). On day 5 post-activation, the KO efficiency was high with up to 85% of downregulation of the TCR expression under all experimental conditions (Figure 1B). The KI efficiency was measured by the expression of Thy1.1 which was 6.9% without any inhibitor treatment (Figure 1C). All tested inhibitors increased the KI efficiency to varying degrees, although it did not reach significance for M3814. CC-115 and PI-103 showed a dose-dependent effect having the strongest increase at 2 µM with a Thy1.1 expression of 9.6% and 10.8%, respectively. Samotolisib led to the overall highest KI efficiency by almost doubling it to 11.6%.

**Figure 1.**
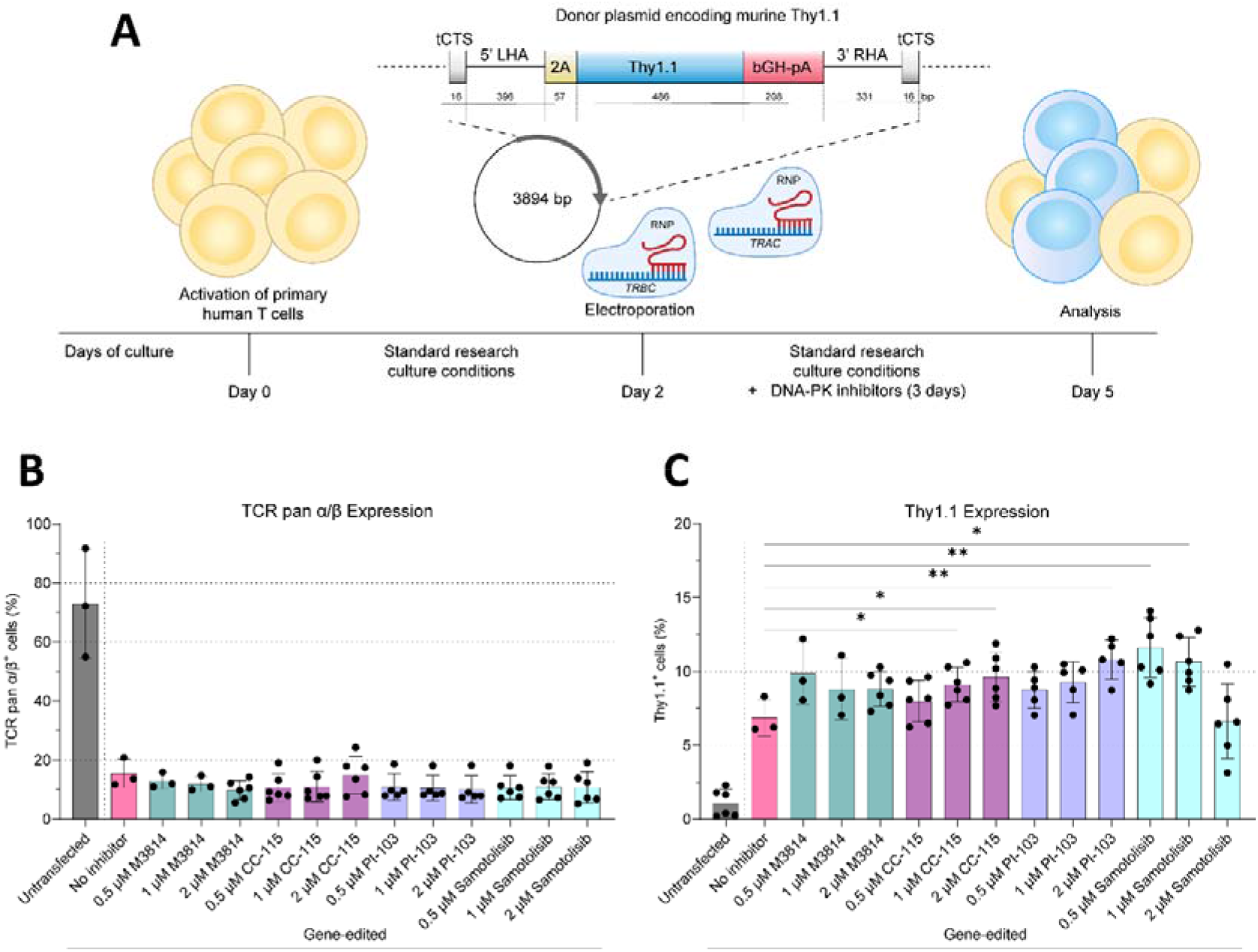
Impact of DNA-PK inhibitors on knockout and knockin efficiency in T cells cultured under standard research conditions. **(A)** Scheme of the experimental design. **(B)** Expression of TCR pan α/β measured by flow-cytometry on day 5 of culture, 3 days after electroporation. **(C)** Expression of Thy1.1 measured by flow cytometry on day 5 of culture, 3 days after electroporation. Both graphs N≥3, each dot represents an individual donor, unpaired *t*-test. tCTS – truncated Cas9 target sequence, LHA – left homology arm, bGH-pA – bovine growth hormone polyadenylation signal, RHA – right homology arm, bp – base pairs.

To investigate whether these inhibitors can be used to increase the KI efficiency in a clinically relevant setting, we adapted our experimental design to generate a GMP-compatible production process of engineered TCR-T cells. In this optimized approach, T cells were gene-edited by CRISPR/Cas9 to express an HLA-A2 restricted NY-ESO-1_157-165_-specific engineered TCR (A2/NY) (25) and a truncated epidermal growth factor receptor (tEGFR) encoded in a Nanoplasmid^TM^ (Aldevron) at the *TRAC* locus. Immediately upon electroporation, the cells were treated for 5 hours with the four different DNA-PK inhibitors and further cultured in a GMP-compatible medium up to day 9 (Figure 2A). Notably, these experiments were performed using the MaxCyte^®^ ExPERT ATx^®^ electroporation system.

**Figure 2.**
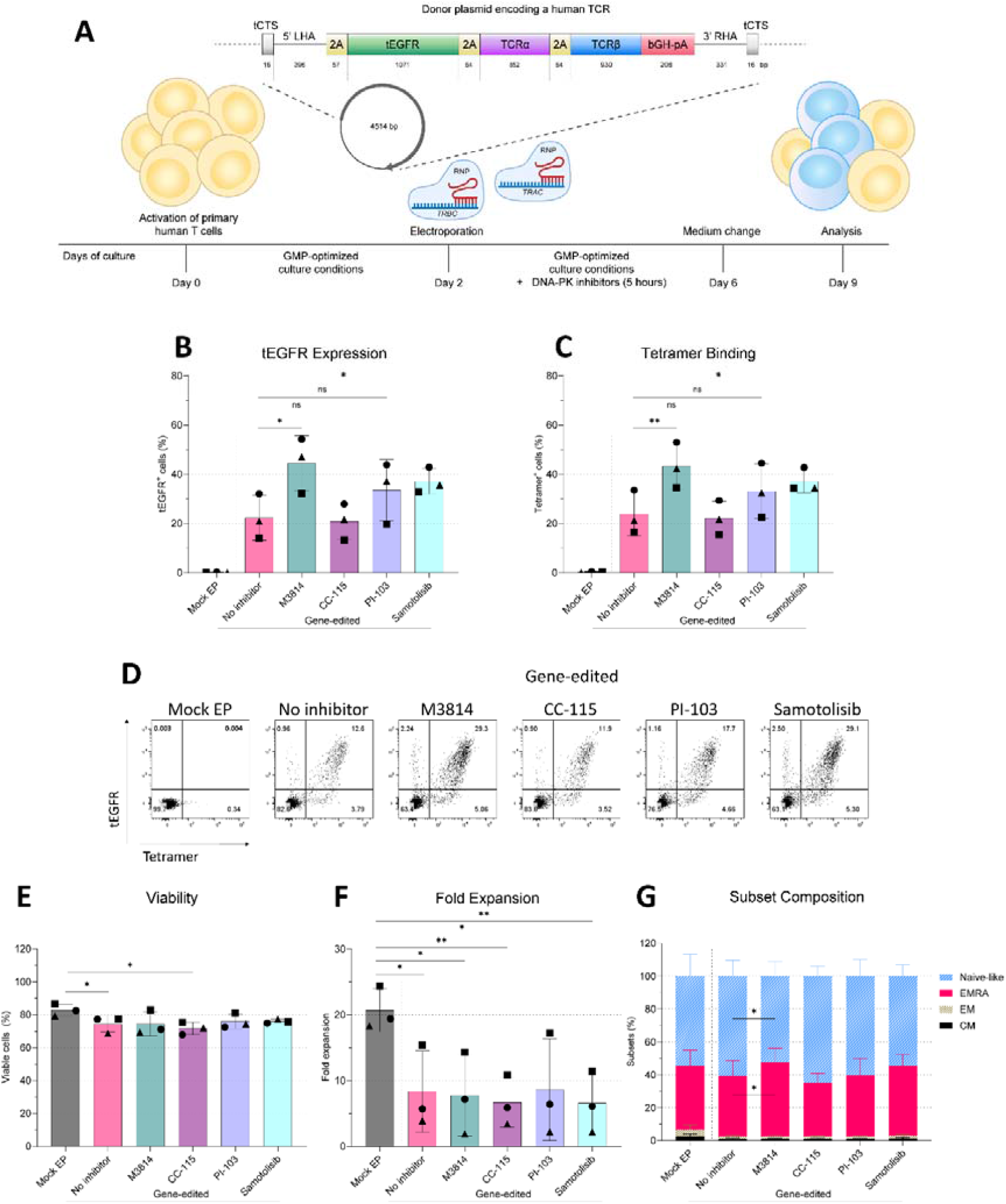
Effect of DNA-PK inhibitors on knockin efficiency, viability, fold expansion and subset composition in T cells cultured under GMP-optimized culture and electroporation conditions. **(A)** Scheme of the experimental design. **(B)** tEGFR expression measured by flow cytometry on day 9 of culture. **(C)** Tetramer binding measured by flow cytometry on day 9 of culture. **(D)** Dotplots of tEGFR expression and tetramer binding of a representative donor from (B) and (C). **(E)** Viability measured by trypan blue staining on day 2 of activation, 5h after DNA-PK inhibitor treatment. **(F)** Fold expansion measured on day 9 of culture in respect to seeding on day 2 post-DNA-PK inhibitor treatment. **(F)** Subset composition measured on day 9 by flow cytometry. All graphs N=3, each symbol for a specific donor, paired *t*-test. tCTS – truncated Cas9 target sequence, LHA – left homology arm, tEGFR – truncated epidermal growth factor receptor, bGH-pA – bovine growth hormone polyadenylation signal, RHA – right homology arm, bp – base pairs, GMP – good manufacturing practice.

With this GMP-compatible experimental design, the KI efficiency was greatly improved, and for a much larger insert (Thy1.1 = 486 bp versus αβTCR+ tEGR = 2853 bp), with an average of 22% tEGFR expression in untreated gene-edited T cells (Figure 2B,D). The treatment with the DNA-PK inhibitors M3814, PI-103 and samotolisib further increased the KI efficiency to an average of 45%, 34% and 38%, respectively. On the other hand, the treatment with CC-115 did not result in an increase of KI efficiency and remained at 21%. Flow cytometric analysis of T cells stained with an A2/NY specific tetramer confirmed KI of the TCR and corresponding to tEGFR expression levels (Figure 2C,D).

Importantly, the DNA-PK inhibition did not adversely impact viability of gene-edited cells, which was on average 74%, five hours after electroporation (Figure 2E). This was overall lower with respect to the mock electroporation (EP) group (83% viability), which reached significance for the experimental groups “no inhibitor” and CC-115. The fold expansion of the mock EP group was on average ∼20-fold while in all gene-edited experimental groups it was ∼10-fold (Figure 2F). At the end of the culture, 55% of cells in the mock EP group had a naïve-like immunophenotype, similar to the gene-edited cells (Figure 2G). The second largest population corresponded to an EMRA phenotype with 38% in the mock EP. DNA-PK inhibitor treatment did not have an effect on the T cell subset phenotype, with the exception of M3814, which resulted in a minor decrease of naïve-like T cells and an increase of the EMRA compartment to 45%. Across all conditions, the CM and EM compartment represented a small fraction, with less than 7% of cells belonging to either subset.

## 4. Discussion

With the aim of augmenting CRISPR/Cas9 KI efficiency by favoring HDR, here we have compared different small molecule DNA-PK inhibitors. We identified that M3814, CC-115, PI-103 and samotolisib increased the KI efficiency to varying degrees in an initial screening experiment. We further investigated the effect of these inhibitors in a modified GMP-optimized process for the generation of A2/NY TCR-T cells co-expressing tEGFR. In these optimized conditions, notably including the use of Nanoplasmid and of the MaxCyte^®^ ExPERT ATx^®^ electroporation system, the KI efficiency was greatly improved even without DNA-PK inhibition.

Using our GMP-optimized process we observed that M3814, PI-103 and samotolisib further improved the KI efficiency by ∼50% by day 9 of cell culture, while the DNA-PK inhibitors CC-115 did not have any effect with respect to the control. Importantly, the treatment of T cells with DNA-PK inhibitors M3814, PI-103 and samotolisib did not lower cell viability up to 5h after electroporation. The variable effect of CC-115 on the KI efficiency under standard versus GMP-optimized culture and electroporation conditions could potentially be explained by the different treatment incubation times of 3 days and 5h, respectively. Of note, the engineered TCR in all conditions is under the control of the endogenous *TRAC* promoter and this strategy has previously shown to delay T cell exhaustion and enhance tumor rejection in comparison to conventional virally engineered T cells (10,12). The fold expansion of gene-edited cells was unaffected by the DNA-PK inhibitor treatment but was overall lower in comparison to the mock EP group in all gene-edited conditions. This is likely a consequence of an intrinsic nucleotide sensing and DNA damage response (26).

Additionally, we have identified that the T cells in all conditions remained in a mostly naïve-like phenotype, at over 50% at the end of culture. This is promising since naïve-like cells and stem cell-like memory T cells have been described to lead to prolonged persistence and superior anti-tumor activity (27,28). Only M3814 increased the EMRA compartment at the cost of naïve-like cells to a minor degree. In summary, site-specific gene editing, ease of use, and the flexibility of CRISPR/Cas9 coupled with the treatment with M3814, PI-103 and samotolisib is a feasible approach for the production of GMP-compatible engineered TCR-T cells. Further investigations are necessary to show an advantage over established cellular engineering protocols and to demonstrate the safety of DNA-PK inhibitors in terms of genomic integrity.

## 5. Materials and Methods

### Isolation and culture of T cells in standard culture conditions

Primary human T cells were isolated from the peripheral blood mononuclear cells (PBMCs) of healthy donors obtained from the Insel Hospital Bern with informed consent. In brief, PBMCs were separated by gradient centrifugation by using Lymphoprep (Axonlab) and CD8^+^ T cells were further negatively selected by using the easySEP (Stemcell Technologies) magnetic bead isolation kit following the manufacturer’s instructions. Isolated cells were cultured in RPMI-1640 (Gibco), supplemented with 10% heat-inactivated FBS (Gibco), 100◻U/ml penicillin and 100◻µg/ml streptomycin sulfate (Gibco), 50 IU/mL IL-2 (Glaxo) and activated for 48h with Dynabeads Human T-Activator CD3/CD28 (Invitrogen) in a ratio of 1:2 T cells to beads.

### CRISPR-Cas9 gene editing of T cells cultured and electroporated under standard conditions

Activated T cells were electroporated with the Sonidel Nepagene instrument 48h post-activation with ribonucleoproteins (RNPs) against *TRAC* and *TRBC* and a donor plasmid (pMA-RQ) encoding murine Thy1.1. In brief, sgRNAs (Synthego) targeting *TRAC* (5’-AGAGTCTCTCAGCTGGTACA-3’) and *TRBC (*ATCGTCAGCGCCGAGGCCTG-3’), were resuspended in water at 88.11 µM. Per electroporation reaction, 8 µL of RNP containing 11 µM *TRAC* sgRNA, 11 µM *TRBC* sgRNA and 7.73 µM Cas9 was prepared in water and incubated at room temperature for 15 minutes. The cells were washed and resuspended in OptiMEM (Gibco) as 1 x 10^6^ cells/90 µL and combined with 8 µL of RNP and 10 µg/1 x 10^6^ cells of a donor plasmid with homology arms flanking the *TRAC* sgRNA PAM site and a structure as previously described (29): truncated Cas9 target sequence (tCTS) (16 base pairs (bp)), left homology arm (396◻bp), P2A, Thy1.1 (486 bp), bovine growth hormone polyadenylation (bGH-pA) signal (208 bp), right homology arm (331◻bp). The electroporation was performed in a green cuvette (Nepa Gene) with the Super Electroporator NEPA21 Type II (Nepa Gene) and the following parameters: poring pulse (voltage 300 V, pulse length 1 ms, pulse interval 50 ms, 2 pulses, 10+ polarity); transfer pulse (voltage 20 V, pulse length 50 ms, pulse interval 50 ms, 5 pulses, 40+/- polarity). Upon electroporation, the cells were cultured in 48-well plates (Falcon) in medium without antibiotics supplemented with DNA-PK inhibitors: M3814, CC-115, PI-103, or samotolisib (all from MedChem Express) at indicated concentrations. The cells were cultured until day 3 post-electroporation.

### Isolation and culture of T cells under GMP-optimized conditions

PBMCs from leukaphereses of healthy donors were obtained from the University Hospital in Lausanne (CHUV) with informed consent. CD4^+^ and CD8^+^ cells were positively selected with CliniMACS GMP CD4 and CD8 microbeads (Miltenyi) by using the Lovo spinning membrane filtration system (Fresenius Kabi) and the CliniMACS plus (Miltenyi), following manufactures’ instructions. The isolated cells were cryopreserved in Cryostor CS10 (StemCell). Upon thawing, the cells were cultured in a G-Rex-100M (Wilson Wolf) with PRIME-XV T Cell chemically defined medium (Irvine Scientific) supplemented with 2500 IU/mL IL-7, 400 IU/mL IL-15 and 25 IU/mL IL-21 (all from CellGenix) (complete medium) and activated with MACS GMP T Cell TransAct (Miltenyi) for 48 h.

### CRISPR-Cas9 KI of T cells under GMP-optimized conditions

Activated T cells were electroporated 48h post-activation with RNPs against *TRAC* and *TRBC* and a donor plasmid to integrate a modified NY-ESO-specific αβ-TCR. RNPs against *TRAC* and *TRBC* were prepared separately. In brief, sgRNAs were resuspended in water at 100 µM. Per electroporation reaction, 40 µL of RNP containing 5 µM sgRNA and 2.5 µM Cas9 was prepared in electroporation buffer (MaxCyte) and incubated at room temperature for 20 minutes. The cells were washed and resuspended in electroporation buffer as 1 x 10^5^/µL. Per reaction, 5 x 10^6^ cells were combined with 20 µL of each RNP and 1 µg/1 x 10^6^ cells of and a donor Nanoplasmid (Aldevron) with homology arms flanking the *TRAC* sgRNA PAM site and the following structure as previously described (29): tCTS (16 bp), left homology arm (396◻bp), P2A, tEGFR (1071 bp), T2A, TCR-α (822 bp) , T2A, TCR-β (933 bp), bgh-PolyA signal (112 bp), right homology arm (331◻bp). Thy1.1 (486 bp), bovine growth hormone polyadenylation (bGH-pA) signal (208 bp), 3′ right homology arm (331◻bp). After electroporation, the cells were left untreated or treated by incubating them for 5h in complete medium supplemented with DNA-PK inhibitors: M3814 (2 µM), CC-115 (2 µM), PI-103 (2 µM), or samotolisib (1 µM). At the end of the incubation, the cells were washed, counted with trypan blue exclusion staining (Life Technologies) and seeded in a G-Rex 24 well plate (Wilson Wolf). At day 4 post-electroporation, 70% of the medium was replaced with fresh medium. Cells were further cultured in complete medium until day 7 post-electroporation.

### Flow cytometric analysis

The T cells generated under standard conditions were stained on day 5 of culture with the antibodies Thy1.1-APC (Biolegend) and pan-TCR-α/β-PE (Beckman Coulter). The T cells generated under GMP-optimized conditions were stained on day 9 of culture with an NY-ESO-MHC I tetramer conjugated with APC (Peptide & Tetramer Core Facility CHUV) and the antibodies EGFR-PE, CCR7-PE-Cy7 and CD45RA-BV510 (all from Biolegend) to differentiate between T cell subsets: naïve-like (CCR7^+^ CD45RA^+^), central memory (CM) (CCR7^+^ CD45RA^-^), effector memory (EM) (CCR7^-^ CD45RA^-^) and terminally differentiated effector memory cells re-expressing CD45RA (EMRA) (CCR7^-^ CD45RA^+^). Viable cells were selected with the LIVE/DEAD™ Fixable Near-IR Dead Cell Stain Kit (Invitrogen). Acquisition was conducted on a BD LSR II (BD Biosciences) and analyzed by using FlowJo software (BD Biosciences).

### Fold expansion analysis

On day 9 of culture, the cells were counted with by trypan blue exclusion staining. The fold expansion was calculated in respect to the seeding on day 2 post-DNA-PK inhibitor treatment.

### Statistical analysis

GraphPad Prism 10 (GraphPad Software) was used for performing statistical analysis. The levels of statistical significance are: *p◻≤◻0.05, **p◻≤◻0.01, ***p◻≤◻0.001 and ****p◻≤◻0.0001.

## Data availability statement

Data will be made available upon request.

## Acknowledgments

We thank our collaborators at MaxCyte for their support in optimizing electroporation conditions for efficient CRISPR/Cas9 knockout/knockin and T cell viability, as well as for their provision of consumables. We also thank the Marson (UCSF, San Francisco, USA) and Corn (ETH, Zurich Switzerland) labs for their technical tips for CRISPR/Cas9 knockin early in this study. We thank and acknowledge the financial support for this work from Ludwig Cancer Research, the University of Lausanne, the ISREC Foundation, the Fondazione Teofilo Rossi di Montelera e di Premuda and the Swiss National Science Foundation (SNF 310030_204326 to M.I.).

## Author contributions

E.D. analyzed data, drafted the manuscript and prepared the figures. G.G.A. and P.R. set-up the CRISPR/Cas9 based KI strategy which was further optimized by MaxCyte electroporation and early inhibitor removal by M.H, all under the supervision of MI. Experiments were performed by M.H., G.G.A., P.R. and S.G., with additional screening experiments by S.G. supervised by K.S. Scientific advice and suggestions were provided by G.C. The overall study was directed and the manuscript edited by M.I. and B.G.

## Glossary

ACT: Adoptive cell transfer
bGH-pA: Bovine growth hormone polyadenylation signal
CAR: Chimeric antigen receptor
CM: Central memory
CRISPR: Clustered regularly interspaced short palindromic repeats
DNA-PK: DNA-dependent protein kinase
EM: Effector memory
EMRA: Effector memory cells re-expressing CD45RA
GMP: Good manufacturing practice
HDR: Homology-directed repair
KI: Knockin
LHA: Left homology arm
NHEJ: Non-homologous end joining
NY-ESO-1: New York esophageal squamous cell carcinoma 1
PBMCs: Peripheral blood mononuclear cells
RHA: Right homology arm
TCR: T cell receptor
tCTS: Truncated Cas9 target sequence
tEGFR: Truncated epidermal growth factor receptor
Thy1.1: Thymocyte differentiation antigen 1.1
TIL: Tumor infiltrating lymphocyte
TRAC: TCR α chain
TRBC: TCR β chain

## References

1. Arienti, F., Belli, F., Rivoltini, L., Gambacorti-Passerini, C., Furlan, L., Mascheroni, L., Prada, A., Rizzi, M., Marchesi, E., Vaglini, M., Parmiani, G., and Cascinelli, N. (1993) Adoptive immunotherapy of advanced melanoma patients with interleukin-2 (IL-2) and tumor-infiltrating lymphocytes selected in vitro with low doses of IL-2. Cancer Immunology, Immunotherapy 36, 315–322

2. Rosenberg, S. A., Yannelli, J. R., Yang, J. C., Topalian, S. L., Schwartzentruber, D. J., Weber, J. S., Parkinson, D. R., Seipp, C. A., Einhorn, J. H., and White, D. E. (1994) Treatment of Patients With Metastatic Melanoma With Autologous Tumor-Infiltrating Lymphocytes and Interleukin 2. JNCI: Journal of the National Cancer Institute 86, 1159–1166

3. Yee, C., Thompson, J. A., Byrd, D., Riddell, S. R., Roche, P., Celis, E., and Greenberg, P. D. (2002) Adoptive T cell therapy using antigen-specific CD8+ T cell clones for the treatment of patients with metastatic melanoma: In vivo persistence, migration, and antitumor effect of transferred T cells. Proceedings of the National Academy of Sciences 99, 16168–16173

4. Rohaan, M. W., Borch, T. H., van den Berg, J. H., Met, Ö., Kessels, R., Geukes Foppen, M.H., Stoltenborg Granhøj, J., Nuijen, B., Nijenhuis, C., Jedema, I., van Zon, M., Scheij, S., Beijnen, J. H., Hansen, M., Voermans, C., Noringriis, I. M., Monberg, T. J., Holmstroem, R. B., Wever, L. D. V., van Dijk, M., Grijpink-Ongering, L. G., Valkenet, L. H. M., Torres, A. A., Karger, M., Borgers, J. S. W., ten Ham, R. M. T., Retèl, V. P., van Harten, W. H., Lalezari, F., van Tinteren, H., van der Veldt, A. A. M., Hospers, G. A. P., Stevense-den Boer, M. A. M., Suijkerbuijk, K. P. M., Aarts, M. J. B., Piersma, D., van den Eertwegh, A. J. M., de Groot, J.-W. B., Vreugdenhil, G., Kapiteijn, E., Boers-Sonderen, M. J., Fiets, W. E., van den Berkmortel, F. W. P. J., Ellebaek, E., Hölmich, L. R., van Akkooi, A. C. J., van Houdt, W. J., Wouters, M. W. J. M., van Thienen, J. V., Blank, C. U., Meerveld-Eggink, A., Klobuch, S., Wilgenhof, S., Schumacher, T. N., Donia, M., Svane, I. M., and Haanen, J. B. A. G. (2022) Tumor-Infiltrating Lymphocyte Therapy or Ipilimumab in Advanced Melanoma. New England Journal of Medicine 387, 2113–2125

5. Irving, M., Lanitis, E., Migliorini, D., Ivics, Z., and Guedan, S.A.-O. Choosing the Right Tool for Genetic Engineering: Clinical Lessons from Chimeric Antigen Receptor-T Cells.

6. Giordano Attianese, G. M. P., Ash, S., and Irving, M. (2023) Coengineering specificity, safety, and function into T cells for cancer immunotherapy. Immunol Rev

7. Irving, M., Lanitis, E., Migliorini, D., Ivics, Z., and Guedan, S. (2021) Choosing the Right Tool for Genetic Engineering: Clinical Lessons from Chimeric Antigen Receptor-T Cells. Hum Gene Ther 32, 1044–1058

8. Foy, S. P., Jacoby, K., Bota, D. A., Hunter, T., Pan, Z., Stawiski, E., Ma, Y., Lu, W., Peng, S., Wang, C. L., Yuen, B., Dalmas, O., Heeringa, K., Sennino, B., Conroy, A., Bethune, M. T., Mende, I., White, W., Kukreja, M., Gunturu, S., Humphrey, E., Hussaini, A., An, D., Litterman, A. J., Quach, B. B., Ng, A. H. C., Lu, Y., Smith, C., Campbell, K. M., Anaya, D., Skrdlant, L., Huang, E. Y., Mendoza, V., Mathur, J., Dengler, L., Purandare, B., Moot, R., Yi, M. C., Funke, R., Sibley, A., Stallings-Schmitt, T., Oh, D. Y., Chmielowski, B., Abedi, M., Yuan, Y., Sosman, J. A., Lee, S. M., Schoenfeld, A. J., Baltimore, D., Heath, J. R., Franzusoff, A., Ribas, A., Rao, A. V., and Mandl, S. J. (2023) Non-viral precision T cell receptor replacement for personalized cell therapy. Nature 615, 687–696

9. Schober, K., Müller, T. R., Gökmen, F., Grassmann, S., Effenberger, M., Poltorak, M., Stemberger, C., Schumann, K., Roth, T. L., Marson, A., and Busch, D. H. (2019) Orthotopic replacement of T-cell receptor alpha- and beta-chains with preservation of near-physiological T-cell function. Nat Biomed Eng 3, 974–984

10. Roth, T. L., Puig-Saus, C., Yu, R., Shifrut, E., Carnevale, J., Li, P. J., Hiatt, J., Saco, J., Krystofinski, P., Li, H., Tobin, V., Nguyen, D. N., Lee, M. R., Putnam, A. L., Ferris, L., Chen, J. W., Schickel, J. N., Pellerin, L., Carmody, D., Alkorta-Aranburu, G., Del Gaudio, D., Matsumoto, H., Morell, M., Mao, Y., Cho, M., Quadros, R. M., Gurumurthy, C. B., Smith, B., Haugwitz, M., Hughes, S. H., Weissman, J. S., Schumann, K., Esensten, J. H., May, A. P., Ashworth, A., Kupfer, G. M., Greeley, S. W., Bacchetta, R., Meffre, E., Roncarolo, M. G., Romberg, N., Herold, K. C., Ribas, A., Leonetti, M. D., and Marson, A. (2018) Reprogramming human T cell function and specificity with non-viral genome targeting. Nature 559, 405–409

11. Oh, S. A., Senger, K., Madireddi, S., Akhmetzyanova, I., Ishizuka, I. E., Tarighat, S., Lo, J. H., Shaw, D., Haley, B., and Rutz, S. (2022) High-efficiency nonviral CRISPR/Cas9-mediated gene editing of human T cells using plasmid donor DNA. J Exp Med 219

12. Eyquem, J., Mansilla-Soto, J., Giavridis, T., van der Stegen, S. J. C., Hamieh, M., Cunanan, K. M., Odak, A., Gönen, M., Sadelain, M., Eyquem, J., Mansilla-Soto, J., Giavridis, T., van der Stegen, S. J. C., Hamieh, M., Cunanan, K. M., Odak, A., Gönen, M., and Sadelain, M. (2017) Targeting a CAR to the TRAC locus with CRISPR/Cas9 enhances tumour rejection. Nature 2017 543:7643 543

13. Müller, T. R., Jarosch, S., Hammel, M., Leube, J., Grassmann, S., Bernard, B., Effenberger, M., Andrä, I., Chaudhry, M. Z., Käuferle, T., Malo, A., Cicin-Sain, L., Steinberger, P., Feuchtinger, T., Protzer, U., Schumann, K., Neuenhahn, M., Schober, K., and Busch, D. H. (2021) Targeted T cell receptor gene editing provides predictable T cell product function for immunotherapy. Cell Reports Medicine 2

14. Camperi, J., Devarajan, S., McKay, A., Tarighat, S., Chen, D., and Hu, Z. (2023) Assessing TCR identity, knock-in efficiency, and potency for individualized TCR-T cell therapy. Journal of Immunological Methods 517

15. Brinkman, E. K., Chen, T., de Haas, M., Holland, H. A., Akhtar, W., and van Steensel, (2018) Kinetics and Fidelity of the Repair of Cas9-Induced Double-Strand DNA Breaks. Molecular Cell 70, 801-813.e806

16. Chu, V. T., Weber, T., Wefers, B., Wurst, W., Sander, S., Rajewsky, K., Kühn, R., Chu, V. T., Weber, T., Wefers, B., Wurst, W., Sander, S., Rajewsky, K., and Kühn, R. (2015) Increasing the efficiency of homology-directed repair for CRISPR-Cas9-induced precise gene editing in mammalian cells. Nature Biotechnology 2015 33:5 33

17. Bozas, A., Beumer, K. J., Trautman, J. K., and Carroll, D. (2009) Genetic Analysis of Zinc-Finger Nuclease-Induced Gene Targeting in Drosophila. Genetics 182

18. Maruyama, T., Dougan, S. K., Truttmann, M., Bilate, A. M., Ingram, J. R., and Ploegh, H. L. (2015) Inhibition of non-homologous end joining increases the efficiency of CRISPR/Cas9-mediated precise genome editing. Nature biotechnology 33

19. Gavande, N. S., VanderVere-Carozza, P. S., Pawelczak, K. S., Mendoza-Munoz, P., Vernon, T. L., Hanakahi, L. A., Summerlin, M., Dynlacht, J. R., Farmer, A. H., Sears, C. R., Nasrallah, N. A., Garrett, J., and Turchi, J. J. (2020) Discovery and development of novel DNA-PK inhibitors by targeting the unique Ku–DNA interaction. Nucleic Acids Research 48

20. Robert, F., Barbeau, M., Éthier, S., Dostie, J., Pelletier, J., Robert, F., Barbeau, M., Éthier, S., Dostie, J., and Pelletier, J. (2015) Pharmacological inhibition of DNA-PK stimulates Cas9-mediated genome editing. Genome Medicine 2015 7:1 7

21. Riesenberg, S., Maricic, T., Riesenberg, S., and Maricic, T. (2018) Targeting repair pathways with small molecules increases precise genome editing in pluripotent stem cells. Nature Communications 9

22. Fu, J., Fu, Y.-W., Zhao, J.-J., Yang, Z.-X., Li, S.-A., Li, G.-H., Quan, Z.-J., Zhang, F., Zhang, J.-P., Zhang, X.-B., Sun, C.-K., Fu, J., Fu, Y.-W., Zhao, J.-J., Yang, Z.-X., Li, S.-A., Li, G.-H., Quan, Z.-J., Zhang, F., Zhang, J.-P., Zhang, X.-B., and Sun, C.-K. (2022) Improved and Flexible HDR Editing by Targeting Introns in iPSCs. Stem Cell Reviews and Reports 18

23. Shy, B. R., Vykunta, V. S., Ha, A., Talbot, A., Roth, T. L., Nguyen, D. N., Pfeifer, W. G., Chen, Y. Y., Blaeschke, F., Shifrut, E., Vedova, S., Mamedov, M. R., Chung, J.-Y. J., Li, H., Yu, R., Wu, D., Wolf, J., Martin, T. G., Castro, C. E., Ye, L., Esensten, J. H., Eyquem, J., and Marson, A. (2022) High-yield genome engineering in primary cells using a hybrid ssDNA repair template and small-molecule cocktails. Nature Biotechnology 41, 521–531

24. Kath, J., Du, W., Pruene, A., Braun, T., Thommandru, B., Turk, R., Sturgeon, M. L., Kurgan, G. L., Amini, L., Stein, M., Zittel, T., Martini, S., Ostendorf, L., Wilhelm, A., Akyüz, L., Rehm, A., Höpken, U. E., Pruß, A., Künkele, A., Jacobi, A. M., Volk, H.-D., Schmueck-Henneresse, M., Stripecke, R., Reinke, P., and Wagner, D. L. (2022) Pharmacological interventions enhance virus-free generation of TRAC-replaced CAR T cells. Molecular Therapy - Methods & Clinical Development 25, 311–330

25. Schmid, D. A., Irving, M. B., Posevitz, V., Hebeisen, M., Posevitz-Fejfar, A., Sarria, J. C., Gomez-Eerland, R., Thome, M., Schumacher, T. N., Romero, P., Speiser, D. E., Zoete, V., Michielin, O., and Rufer, N. (2010) Evidence for a TCR affinity threshold delimiting maximal CD8 T cell function. Journal of immunology 184, 4936–4946

26. An, J., Zhang, C. P., Qiu, H. Y., Zhang, H. X., Chen, Q. B., Zhang, Y. M., Lei, X. L., Zhang, C. X., Yin, H., and Zhang, Y. (2024) Enhancement of the viability of T cells electroporated with DNA via osmotic dampening of the DNA-sensing cGAS-STING pathway. Nature Biomedical Engineering 8, 149–164

27. Hinrichs, C. S., Borman, Z. A., Cassard, L., Gattinoni, L., Spolski, R., Yu, Z., Sanchez-Perez, L., Muranski, P., Kern, S. J., Logun, C., Palmer, D. C., Ji, Y., Reger, R. N., Leonard, W. J., Danner, R. L., Rosenberg, S. A., and Restifo, N. P. (2009) Adoptively transferred effector cells derived from naïve rather than central memory CD8+ T cells mediate superior antitumor immunity. Proceedings of the National Academy of Sciences 106, 17469–17474

28. Gattinoni, L., Lugli, E., Ji, Y., Pos, Z., Paulos, C. M., Quigley, M. F., Almeida, J. R., Gostick, E., Yu, Z., Carpenito, C., Wang, E., Douek, D. C., Price, D. A., June, C. H., Marincola, F. M., Roederer, M., and Restifo, N. P. (2011) A human memory T cell subset with stem cell-like properties. Nat Med 17, 1290–1297

29. Nguyen, D. N., Roth, T. L., Li, P. J., Chen, P. A., Apathy, R., Mamedov, M. R., Vo, L. T., Tobin, V. R., Goodman, D., Shifrut, E., Bluestone, J. A., Puck, J. M., Szoka, F. C., and Marson, A. (2020) Polymer-stabilized Cas9 nanoparticles and modified repair templates increase genome editing efficiency. Nat Biotechnol 38, 44–49

